# Defining the genetic determinants of CD8^+^ T cell receptor repertoire in the context of immune checkpoint blockade

**DOI:** 10.1101/2024.11.04.618564

**Authors:** Esther S. Ng, Orion Tong, Chelsea Taylor, Robert Watson, Bo Sun, Gusztav Milotay, Sophie MacKay, James J Gilchrist, Martin Little, Benjamin P Fairfax, Yang Luo

## Abstract

CD8^+^ T cells play a central role in the cancer response to immune checkpoint blockade (ICB) treatment, with activity predicated upon antigen recognition by the associated T cell receptor (TCR) repertoire. The contribution of genetic variation to this in cancer treatment is under-explored. We have conducted a genome-wide and human leukocyte antigen (HLA)-focused analysis of CD8^+^ T cell TCR repertoire to identify genetic determinants of variable gene (V-gene) and CDR3 K-mer usage from samples taken prior to and after ICB treatment (n=250). We find 11 genome-wide significant *cis* associations and 10 *trans* associations, primarily to the HLA, with V-gene usage meeting permuted P-value thresholds. Notably, TCR clones containing V-genes associated with HLA were less likely to be persistent across treatment. In a single-cell experiment, we find cells with HLA-matched TCR clones have increased tumor reactivity expression profiles and patients with HLA-matched TCR clones have improved overall survival. Our work indicates a complex relationship between genotype and TCR repertoire in the context of treatment with ICB, which has novel implications for understanding determinants of treatment response and patient outcomes.

**One Sentence Summary:** TCR repertoire is strongly associated with specific HLA alleles in cancer patients, but immune checkpoint blockade influences this association.

## INTRODUCTION

Immune checkpoint proteins, of which the cell-surface receptors programmed cell death protein 1 (PD-1) and cytotoxic T-lymphocyte-associated antigen 4 (CTLA-4) are archetypal examples, play key roles in negatively regulating T cell responses, limiting deleterious inflammation and autoimmunity. In the context of cancer, induction and ligation of PD-1 and CTLA-4 are implicated in the development of T-cell exhaustion and suppression of anti-tumor immunity. Correspondingly, treatment with immune checkpoint blockade (ICB) leads to re-invigoration of exhausted T cells, increasing their proliferation, cytotoxicity and broadening the TCR repertoire[1, 2, 3]. Treatment with ICB has improved outcomes across numerous cancers, most notably metastatic melanoma (MM) where treatment with combination ICB to CTLA-4 (ipilimumab) and PD-1 (nivolumab) for MM is associated with extending median survival from months to beyond five years [4, 5].

CD8^+^ T cells are central to T cell mediated anti-tumor responses to both neoantigens and Tumour Associated Antigens (TAA) presented by Class I Human Leukocyte Antigen (HLA) alleles in complex with *β*-2 microglobulin. Successful recognition of HLA presented antigens utilises TCR incorporating *α* and *β* chains encoded on chromosome 14 and 9 respectively. Within the TCR *β* chain the first and second complementarity-determining regions (CDR1, CDR2) primarily interface with HLA *α* helices, whilst interactions with the hypervariable CDR3 region are the critical determinant of peptide recognition [6]. As such, in both the *α* and *β* chains, the CDR3 loops have the highest sequence diversity and are the principal determinants of receptor specificity [7]. The CDR1 and 2 loops are determined by the V-gene segment in the TCR whilst CDR3 hypervariability is secondary to V(D)J recombination during early T cell maturation. The process of gene segment recombination to encode a final full length transcript results in a combinatorial possibility of 10^10^ possible TCR sequences [8], vital for enabling recognition of a vast array of self or pathogen derived peptides.

At the individual level the TCR repertoire is determined by a complex interplay between antigen exposure and HLA determined propensity to present specific peptides. Early thymic selection is crucial in repertoire determination, with deletion from the repertoire of T cell clones expressing TCRs either unresponsive or overtly reactive to self-peptide MHC complexes[9, 10]. The interaction between germline genetics at the MHC and TCR repertoire thus has significant implications on our understanding of the pathophysiology of directly immune-mediated diseases and those where immunity plays a key role in clinical responses to therapeutics including cancer. However, in the context of ICB therapy, the relationship between germline genetics, including classical HLA alleles, and TCR usage, as well as any association to clinical response to treatment are largely unexplored.

Here we have investigated the association between genetic factors and CD8^+^ TCR repertoire composition across 250 patients receiving ICB for cancer, integrating germline genotyping and TCR sequencing both prior to and post ICB treatment. We describe a complex relationship between genotype determined Class I HLA allele carriage and the TCR repertoire, specifically identifying a number of HLA-matched TCR chains. By exploring single cell RNA sequencing (scRNA-seq) data from patients receiving ICB we find CD8^+^ T cell clones incorporating HLA-matched TCR demonstrate increased expression of tumor reactivity associated genes. Finally, we find that presence of HLA-matched TCR clones is associated with overall survival in patients receiving ICB for MM.

## RESULTS

### Exploring CD8^+^ T cell receptor repertoire

We performed RNA sequencing of peripheral CD8^+^ T cells isolated prior to the first and second cycles of ICB treatment from 250 patients receiving ICB for cancer (**Supplementary Table 1**). By correlating *α* and *β* chain V-gene usage between patients we identified conserved patterns of V-gene repertoire frequency **(Figure 1A,B)**. Principal component (PC) analysis of correlations between *α* and *β* chain V-genes demonstrated further distinct relationships in chain usage between *α* and *β* chains (**Supplementary Figure 1A,B**). Notably, the first PC of *α* chain V-gene usage was dominated by *TRAV1-2*, which was markedly separate from other chains, and forms the key TCR for MR1 ligated Mucosal-associated invariant T cells (MAIT) cells [11]. *TRAV12-2* formed one of outlier chains for the second PC, clustering relatively close to *TRAV1-2* - interestingly *TRAV12-2* has been described to be used by atypical MR1 reacting MAIT cells [12]. Correspondingly, a reciprocal pattern was observed in the first PC of the *β* chain usage which was dominated by *TRBV6-4*, the key partner of *TRAV1-2* in MAIT cells - thus it would appear that whilst there is limited structure within paired CD8^+^ TCR chain usage, reactivity to MR1 versus HLA is the leading driver of variation.

**Figure 1.**
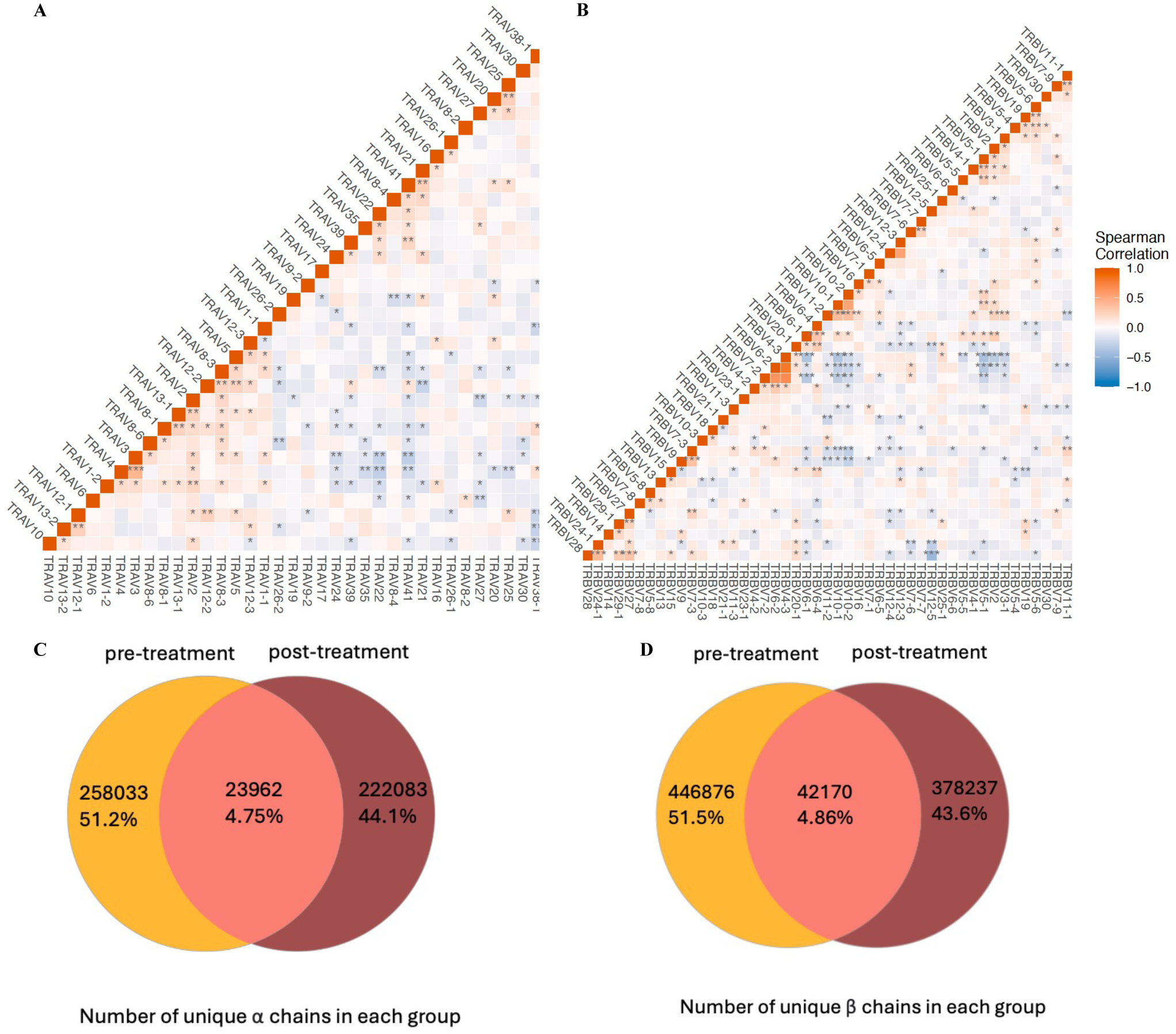
Summary of TCR chains in our study. Spearman correlation between each TCR (**A**) *a* (**B**) TCR *β* V-gene usage in our cohort. In each plot, V-gene usage is calculated by normalising the number of unique clones per individual. \**p <* 0.05; \*\**p <* 0.01.(**C**) Number of TCR *a* chains and (**D**) TCR *β* chains in three groups of cells - those seen only in individuals pre-ICB treatment, those seen in individuals only post-ICB reatment and those seen both pre- and post-ICB treatment.

For 179 patients we had paired pre and post ICB TCR data. In these we identified 504,078 unique *α*-chain clones and 867,283 unique *β*-chain clones. Analysis of chain conservation across ICB treatment demonstrated that only a small number of clones were resampled across both time points, likely reflecting the relatively small proportion of total TCR being sampled as well as ICB induced T cell mitosis **(Figure 1C,D)** [13].

### GWAS of V-gene usage reveals *cis*- and *trans*-associations

We proceeded to explore the relationship between genetics and TCR usage. No difference in overall V-gene usage pre- and post-treatment was observed in PCA (**Supplementary Figure 2**), and henceforth focused our association analysis on pre-treatment samples where our power was greatest to detect effect (n=250). Genome-wide genotyping was performed across samples, resulting in 486,469 SNPs available for post-QC analysis. We defined V-gene usage for *α* and *β* chains as normalised count of unique clones per V-gene and modelled this as a quantitative trait. For each V-gene, we performed additive linear regression to identify genetic variants associated with V-gene usage, correcting for the first two genetic PCs, two TCR PCs, age, gender and cancer type (**Supplementary Figures 3,4**). To account for multiple testing and also correlation between V-gene usage, we permuted the phenotype dataset 1,000 times, preserving individual level V-genes but re-ordering the samples, resulting in a permutation P-value threshold of 3.83 x 10^−9^ for *β* chain V-gene (TRBV) and 3.46 x 10^−9^ for *α* chain V-gene (TRAV, **Supplementary Figure 5A,B**). In total, we performed genome-wide association studies for 47 *β* chains and 42 *α* chains V-genes from pre-ICB treatment samples. After running *>* 40 million regression models we observed, as per previous observations [14], strong *cis* associations to a subset of the *α* and *β* chains, corresponding to their respective genomic loci (**Figure 2A**, **Figure 2B**). Additionally we noted a strong secondary *trans*-acting signal to both chains corresponding to the MHC region at 6p23.

**Figure 2.**
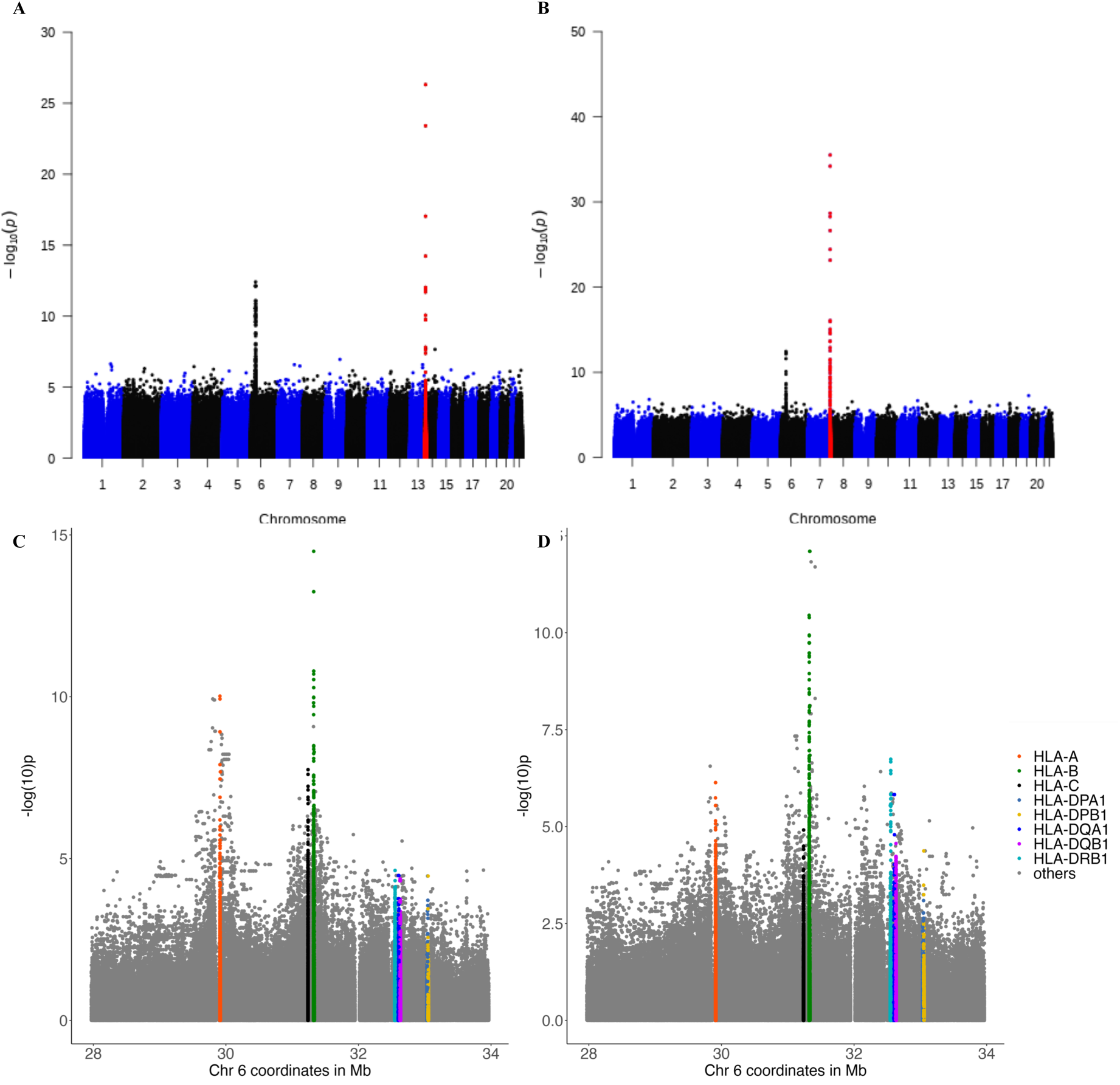
Association between TCR V-gene usage and germline genetic variation. (**A**) Manhattan plot of association between TCR *a* chain V-gene usage. The TRAV gene locus is highlighted in red. (**B**) Manhattan plot of TCR *β* chain V-gene usage. The TRBV gene locus is highlighted in red. The strength of association is indicated by the *—* log_10_ of the P-value of the linear model fitted between each V-gene usage and each variant on the y-axis. The x-axis shows the genomic position (in GRCh38). All V-genes are plotted together and TRAV and TRAV gene loci are highlighted in red. Locus plot of association between (**C**) TRAV and (**D**) TRBV usage and variants in the MHC region. Classical HLA genes are annotated with different colours in the legend. (**E**) Heatmap of omnibus test of each HLA amino acid position against V-gene usage.

The mechanisms whereby *cis*-acting polymorphisms may influence V-gene usage may be similar to standard expression quantitative loci, with polymorphisms in regulatory regions such as promoters and enhancers regulating expression [15, 16]. However, by providing equal weighting to clones irrespective of numbers of copies detected, our approach gives insights into propensity for clonal formation. Thus *cis* acting variants are more likely to influence propensity to recombination, defined by the local three dimensional structure of the loci [17, 18]. In total we found clones containing nine different TRBV-genes were associated with variants in *cis* at P-values below the permuted significance threshold (**Supplementary Table 2**). The most significant association was for *TRBV28* was with rs4726571, 3.7Kb upstream (*β* = *—*0.957, *P* = 2.12 *×* 10*^—^*^40^). At the *α* locus only *TRAV26-2* and *TRAV38-1* demonstrated associations with clonal usage after correcting for multiple testing, the most significant being with rs1023437 which is 41Kb downstream from *TRAV26-2* (*β* = *—*1.035, *P* = 4.88 *×* 10*^—^*^27^). We performed further conditional analysis on all the significant V-genes, controlling for primary *cis* signals. Only one secondary signal was observed to *TRBV4-3*, with the lead SNP being rs361489 (*β* = 0.571, *P* = 9.83 *×* 10*^—^*^12^) and peak secondary signal to rs17249 (*β* = *—*0.460, *P* = 2.66 *×* 10*^—^*^11^), indicating multiple germline variants determining the usage of this V-gene.

### V-gene usage is associated with classical HLA alleles

We observed a *trans* locus associated with both TRAV and TRBV genes that maps to 6p23, which corresponds to the MHC region. To further define the associations here we associated V-gene usage with imputed SNPs in the MHC region as well as classical HLA alleles and coding polymorphisms. In total we tested 16,781 SNPs, 174 classical alleles, and 2,119 amino acids. To control for false positives, we performed a permutation analysis by randomly shuffling the phenotypes and fitting the regression model 1,000 times deriving empirical P-value thresholds of 3.37 x 10^−7^ for *β* chain and 3.61 x 10^−7^ for *α* chain V-genes respectively (**Supplementary Figure 5C,D**). In total, we observed five *β* chain V-genes and five *α* chain V-genes that had significant associations with variants within the MHC region (**Figure 2C,D**, **Supplementary Table 3**, **Supplementary Figure 6-8**).

We found that the most significant association in the *α* chain was HLA-A*02 with *TRAV12-2* (*β*=0.689, P=3.04 x 10^−13^, **Figure 3A,B**). Interestingly a strong bias towards *TRAV12-2* usage has been described for melanocyte antigen (Melan-A) specific TCR, with predominant interaction between the *TRAV12-2* chain and the HLA-A2/Melan-A peptide located in the CDR1loop (Gln31) [19]. Given Melan-A is one of the most common melanoma associated antigens, these results suggest that germline determined *TRAV12-2* usage may play a role in anti-melanoma immunity. There were no secondary signals in the conditional analysis of any of the TRAV-genes. The strongest association observed to the *β* chain was between *TRBV19* and rs2250287 (*β*=0.760, P=7.93 x 10^−13^, **Figure 3C**) with the top classical HLA allele associated with *TRBV19* being HLA-B*44 (*β*=0.734, P=1.83 x 10^−10^, **Figure 3D**) in linkage disequilibrium with rs2250287 (r^2^=0.882). At four-digit HLA resolution, *TRBV19* was associated with both HLA-B*44:02 (*β*=0.576, P=2.78 x 10^−5^) and HLA-B*44:03 (*β*=0.725, P=1.23 x 10^−4^). Interestingly, HLA-B*44:02 is associated with protection against multiple sclerosis in several studies [20, 21, 22], whilst in the context of melanoma carriage of the B*44 supertype has been associated with improved outcomes to ICB treatment[23] - although this was not reproduced in a follow-up study[24]. Notably though, as per *TRAV12-2*, *TRBV19* is enriched for TCR in T cells reactive to Melan-A in patients with metastatic melanoma [19]. We further performed conditional analysis and observed only one secondary signal among the significantly associated V-genes - *TRBV19* with the top secondary SNP being rs62395263 (*β*=0.705, P=2.96 x 10^−7^).

**Figure 3.**
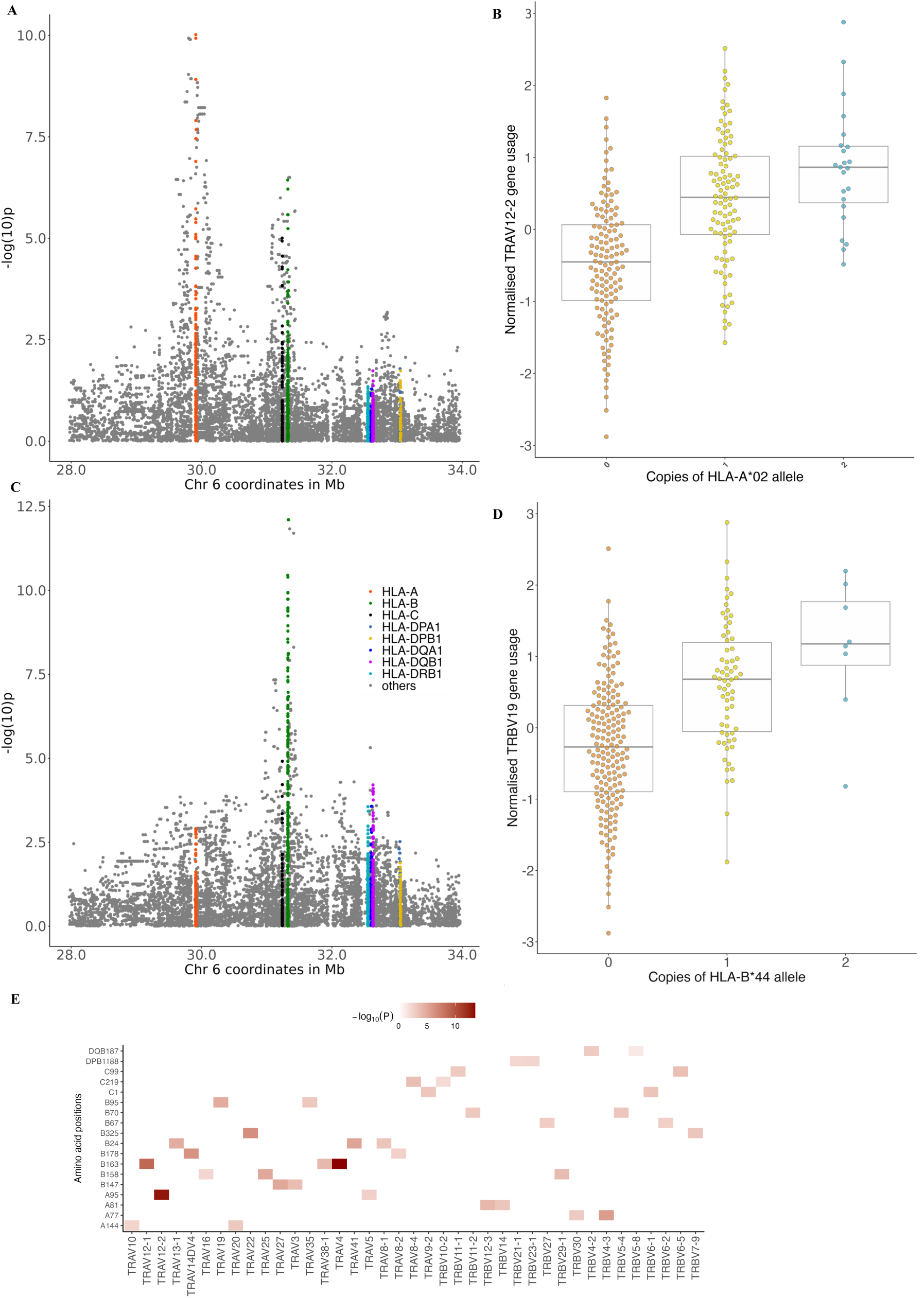
Association between V-gene usage and MHC variants. The MHC locus plot of (**A**) *TRAV12-2* and (**C**) *TRBV19*. The x-axis shows the genomic positions of chromosome 6 (build GrCh38), and the y-axis is the *—* log_10_(P) obtained from two-sided regression analyses. (**B**) *TRAV12-2*usage plotted against dosage of the most significant classical HLA allele association, HLA-A*02. Each dot represents an individual. (**D**) *TRBV19* usage plotted against dosage of the most significant classical HLA allele, HLA-B*44. Box plots show median (horizontal bar), 25th and 75th percentiles (lower and upper bounds of the box, respectively) and 1.5*z*IQR (or minimum and maximum values if they fall within that range; end of whiskers).

To determine whether specific HLA amino acid positions were associated with V-gene usage, we tested variable amino acid positions by grouping haplotypes carrying a specific residue at each position in an additive model (**Supplementary Table 4, Supplementary Figure 9**). We found that the most significant association in *α* chain was between *TRAV12-2* and position 74 (exon2) in HLA-A (*P_ommnibus_* = 2.93 x 10^−13^), whilst the most significant association in *β* chain was between *TRBV19* and position 45 (exon2) in HLA-B (*P_ommnibus_*= 5.08 x 10^−11^). These positions had stronger association signals than any single SNP or classical HLA allele, and fall within the peptide-binding groove of the respective HLA protein, indicating that variation in the amino acid content of the peptide-binding groove is the major genetic determinant of V-gene usage, likely reflecting presentation of a common antigen being the key driver (**Figure 3E**).

### CDR3 amino acid K-mers are associated with HLA alleles

In both *α* and *β* TCR chains, CDR3 loops - which form the hypervariable peptide contacting region - have the greatest diversity in sequence and are the crucial determinants of antigen binding specificity[25]. We hypothesised that specific CDR3 sequences would demonstrate reciprocal association with variants in the MHC region. To test this, we employed a K-mer based approach across TCR mapped from the CD8^+^ T cell bulk sequencing data, using a sliding window of seven amino acids to quantify K-mer motif usage in the CDR3 region (6,654 *β* chain and 2,306 *α* chain K-mers respectively) across individuals. Linear models were subsequently utilised to test the association between CDR3 K-mers and HLA alleles, incorporating the same covariates as prior analyses.

Whilst no significant associations were identified for the *α* chain, we identified two significantly associated K-mers in the *β* chain (**Supplementary Figure 10**). The most significant *β* chain CDR3 K-mer association was between TGDSNQP and HLA-B*35:01 (*β*= 1.41, P=1.57 x 10^−13^) whilst the K-mer TSGDYNE was associated with rs3763288 (*β*= 1.12, P=2.66 x 10^−10^), an intronic variant in *MICA*, encoding MICA (Major Histocompatibility Complex Class I-like molecule A), a ligand for the NK and CD8^+^ T cell activating immunoreceptor NKG2D [26] which triggers a cytotoxic response upon ligation[27]. Cell surface release of MICA in a soluble form (sMICA) has been implicated in cancer cell escape from natural killer cell immune surveillance [28] and rs3763288 forms a quantitative trait locus for blood sMICA levels [29]. To test whether the observed associations with CDR3 K-mers were influenced by V-gene usage, we performed a correlation analysis between CDR3 K-mers and V-gene usage. Whilst no V-genes were associated with TGDSNQP, indicating that this K-mer association is independent of V-gene pairing, TSGDYNE was associated with *TRBV19* (pearson r=0.276, P=9.61 x 10^−6^). We repeated our analysis of *TSGDYNE*against rs3763288 conditioning on *TRBV19* and the association remained significant (*β*=1.01, P=1.06 x 10^−8^), suggesting that whilst TSGDYNE was frequently observed in CDR3 from *TRBV19* containing TCR, the association between TSGDYNE and rs3763288 is not dependent upon *TRBV19*.

### Refinement of genetic signal at single-cell resolution

To validate our associations we performed scRNA-seq with VDJ mapping on T Cells from 59 individuals with melanoma, 55 of whom were also included in the bulk RNA analysis. A further strength of this approach being it enabled examination of cell-type specific associations across T cell subsets (**Supplementary Table 5**).

Testing the lead *β* chain *cis* association, rs4726571 with *TRBV28*, demonstrated replication at the single-cell level across all CD8^+^ cells (*β*=-0.339, P=0.00412) and was also observed in all CD4^+^ cells (*β*=-0.320, P=0.00551), indicating a pan T cell mechanism. Notably, the association was stronger in CD8 terminally differentiated effector memory cells re-expressing CD45RA (TEMRA) cells (*β*=-0.489, P=5.53×10^−5^) and CD8^+^ T Effector Memory (TEM) cells (*β*=-0.426, P=1.35×10^−4^) compared to CD8 Naive cells (*β*=-0.236, P=0.0700), suggesting additional maintenance of clonal abundance due to antigen recognition.

In general, associations in *trans* are harder to replicate due to the typically secondary mechanisms of action. Given this, it was noteworthy that we observed the same direction of effect of the leading MHC*×*V-gene association (**Supplementary Table 6**) between *TRBV19* and HLA-B*44 across all CD8 T^+^ cells (*β*=0.0823, P=0.391). In keeping with this being secondary to antigen presentation via Class I HLA to CD8^+^ cells, we found no evidence of association to CD4^+^ T cells (*β <* 0). Further in keeping with this association being secondary to antigen presentation, the association across CD8^+^ T cell subsets demonstrated a stronger signal in CD8 TEMRA cells (*β*=0.373, P=0.0398) compared to CD8 Naive cells (*β*=0.0119, P=0.950).

### ICB alters HLA V-gene association

Our findings regarding the frequency of V-gene usage and HLA allele status were based on CD8^+^ T cells from patients prior to ICB treatment. We wanted to explore the effect of ICB treatment on potential HLA-V-gene selection. A key strength of the dataset is that most patients had paired CD8^+^ T cell samples before and after their first cycle of ICB treatment, allowing us to explore genotype*×*V-gene usage post-treatment. Focusing on the 179 individuals for which paired samples were available, we sub-divided clones into three groups: those only observed pre-treatment (*Unstable*), those observed post-treatment only (*Novel*), and those present across both samples (*Persistent*). We selected all V-genes that had a significant association with variants in the MHC region (**Supplementary Table 3**), and extracted the top classical HLA alleles with that V-gene. We define a clone to be HLA-matched if it had both the presence of the associated HLA allele and V-gene. We subsequently calculated the proportion of HLA-matched clones for each individual within each clone group. For the *β* chain, we found that the proportion of HLA matched clones in each individual in the *unstable* group was significantly higher (median = 0.238) than in the *novel* group (median = 0.132, P=0.000436, Wilcoxon test) and *persistent* groups (median = 0.125, P=0.000107, Wilcoxon test, **Figure 4A**). Although the median proportion of HLA-matched clones was higher in the novel compared to persistent group, this was not statistically significant. Similarly for the *α* chain the proportion of HLA-matched clones was also higher in the unstable group than the persistent group (P=0.00229) although there was no significant difference in proportions between unstable and novel groups or between persistent and novel groups (**Figure 4B**). The observation that the median proportions of HLA-matched clones is highest in the *unstable* group suggests that, in the context of ICB therapy which acts on exhausted T cells, clones that grow and shrink with treatment - and potentially responding to antigen presentation - are more likely to be HLA-selected (**Supplementary Table 7**). This is illustrated in the leading *β* chain association between *TRBV19* with HLA-B*44 which was strongest in the unstable and novel groups (**Figure 4C-E**).

**Figure 4.**
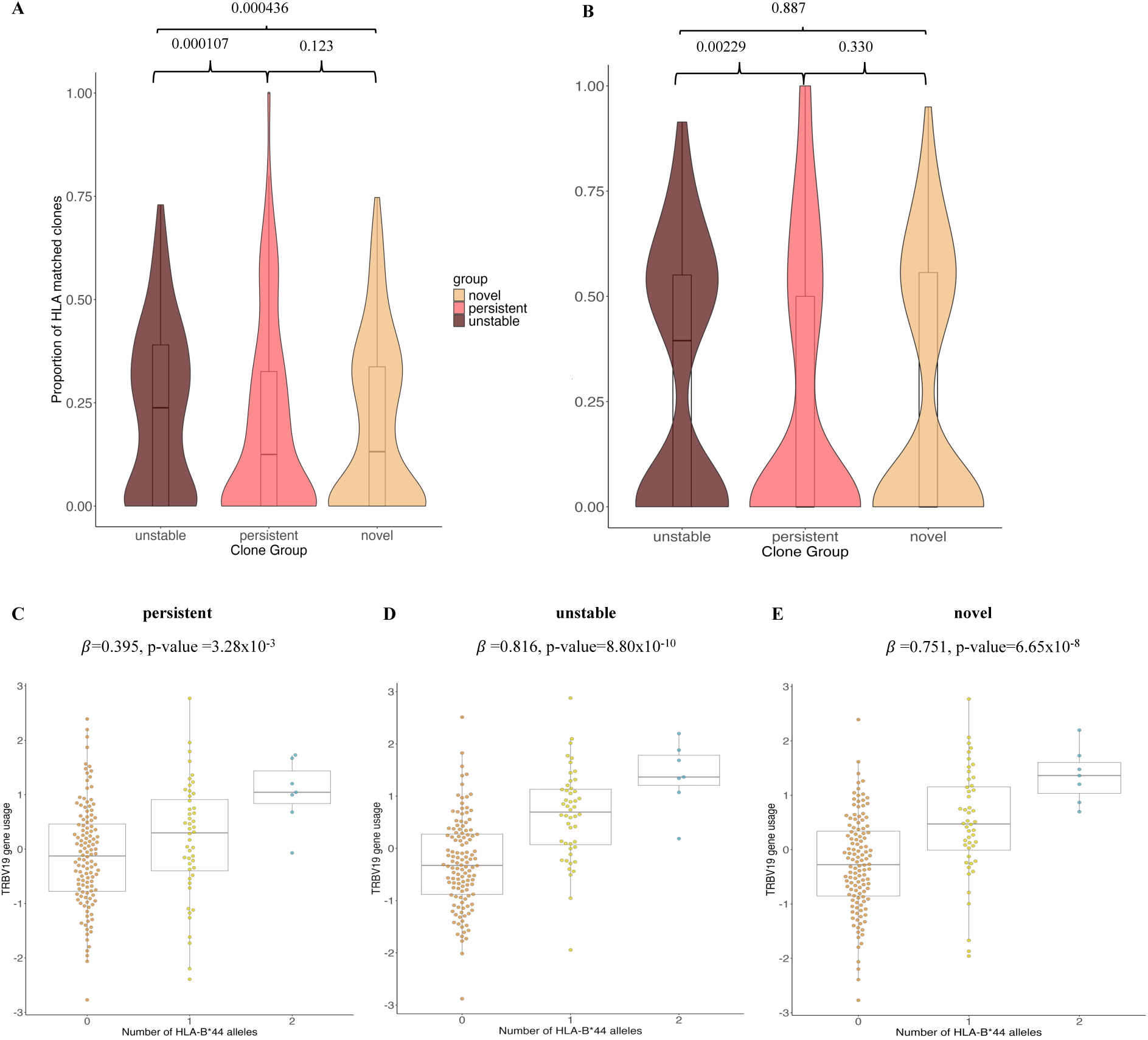
HLA-Vgene association across ICB treatment. The proportion of HLA-matched clones in three groups of clones - those only observed pre-treatment (*Unstable*), those observed post-treatment only (*Novel*), and those present across both samples (*Persistent*) in **A** *β* chain and (**B**) *a* chain. Each dot represents an individual and the paired wilcoxon test P-values are represented above the plots for comparison between the groups.*TRBV19* plotted against HLA-B*44 in (**C**) persistent, (**D**) unstable and (**E**) novel of clones demonstrating differential relationship between V-gene usage and copies of classical HLA alleles.

### Cells with HLA matched V-gene demonstrated increased tumor reactivity expression profiles

Given that unstable clones are more likely to carry TCRs that demonstrate HLA selection, we sought to determine whether such HLA-selected clones showed evidence of tumor reactivity. To do this, we applied a set of 20 genes identified to characterise tumor-reactive cells based on single-cell sequencing in 59 melanoma patients [30], generating a tumor-Reactivity Score (TRS, **Supplementary Table 8**). Using V-genes significantly associated with variants in the MHC region, we extracted the top classical HLA allele associated with each V-gene and dichotomized cells according to whether they carried HLA-matched TCRs or not. For each cell, we calculated its TRS and determined the association between TRS and HLA-matching status using a linear model. We showed that HLA-matched pre-ICB CD8^+^ cells displayed a significantly increased TRS (*β*=0.0870, P=5.82×10^−4^), in keeping with cancer-related antigenic pressure in these patients. We subsequently explored whether the increased TRS seen in HLA-matched cells were also seen in pre-defined cell subtypes (**Figure 5**). Pre-treatment the most robust association was observed in T effector memory cells (CD8 TEM, *β*=0.155, P=7.14×10^−4^, **Supplementary Table 9**). To see if this effect was driven by singlet or expanded clones, we tested this association across both categories, finding that the association between TRS and HLA-matching status was confined to cells from expanded clones (singlets: *β*=0.0415, P=0.256, expanded clones: *β*=0.140, P=6.87 x 10^−5^). Consistent with this observation, we found that TRS was positively correlated with clone size (P=7.21 x 10^−34^, Spearman r = 0.0558).

**Figure 5.**
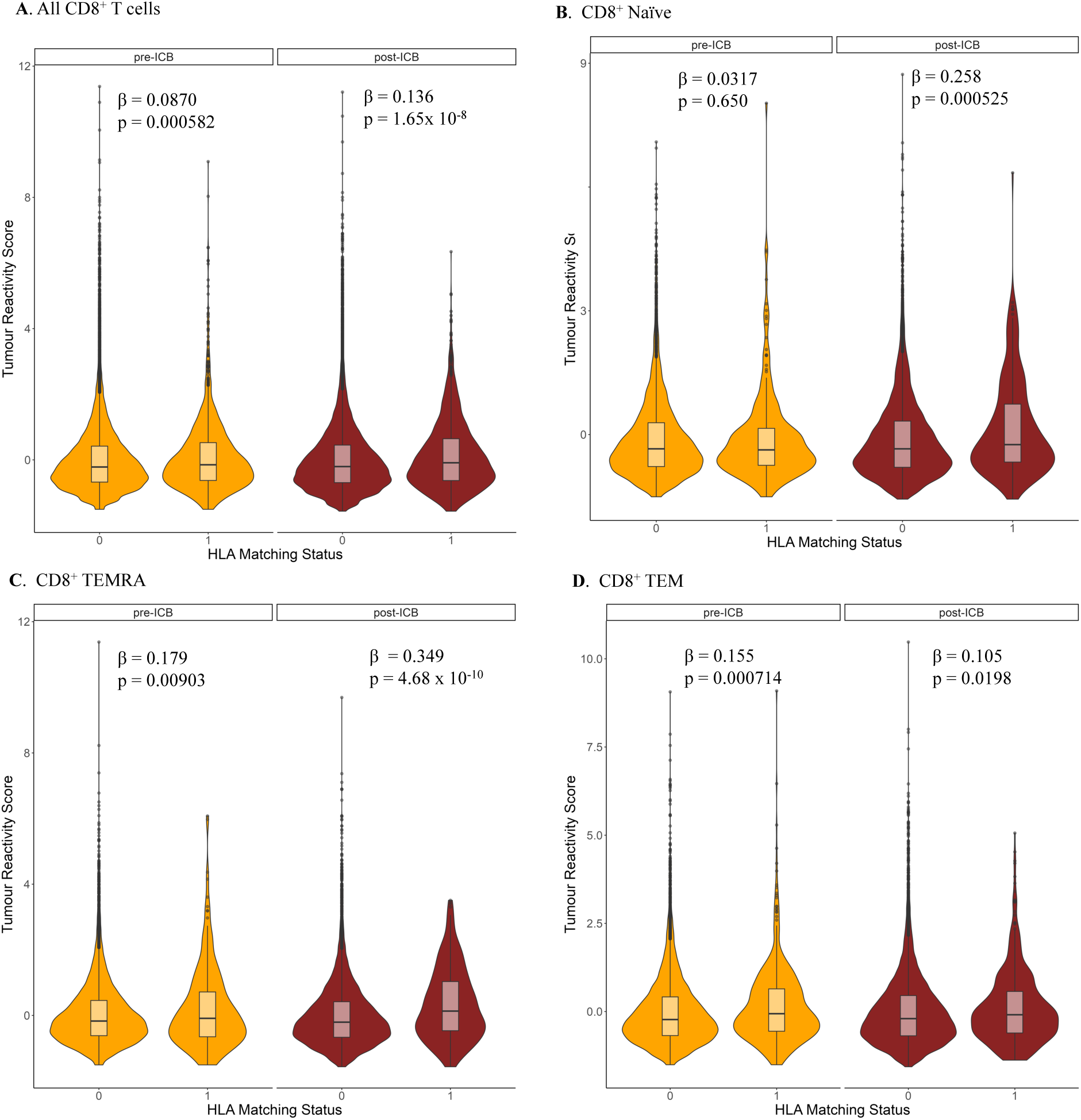
Association between V-gene usage and HLA matching status. (**A**) HLA matching status (0-unmatched, 1-matched) plotted against tumor reactivity score (TRS) for all CD8^+^ cells pre and post treatment with each dot representing one cell. The subplots are annotated with anova P values comparing the model with HLA matching status with the null model without HLA matching status. (**B-D)** In each subplot, tumor reactivity score is plotted against HLA matching status with each dot representing one cell. Each subplot represents a CD8^+^ cell subset, showing that the largest difference occurs within T Effector Memory (TEM) cells before treatment but within terminally differentiated effector memory cells re-expressing CD45RA (TEMRA) cells after treatment.

To investigate whether the observed associations were different in post-treatment samples, we repeated this analysis for all cells from a subset of 56 individuals after the first cycle of ICB treatment. We demonstrated that the association between TRS and HLA matching among all CD8^+^ cells was far stronger post-ICB (*β*=0.136, P=1.65×10^−8^) and was also detectable across multiple cell subsets, with the strongest association in CD8 TEMRA (*β*=0.349, P=4.68×10^−10^), followed by naïve cells (*β*=0.258, P=5.25 *×* 10*^—^*^4^) and CD8 TEM cells (*β*=0.105, P=0.0198). We also separated the cells into whether they were singlets or from expanded clones, observing that post-ICB, both singlets and expanded clones demonstrated increased TRS if HLA-matched (singlets: *β*=0.150, P=3.85 *×* 10*^—^*^5^, expanded: *β*=0.130, P=7.18 *×* 10*^—^*^5^).

We noted that the strongest association between tumor reactivity and the HLA associated V-genes was observed in CD8 TEM cells before ICB treament and in CD8 TEMRA cells after treatment. This shift may be linked to previously described phenomenon of ICB-induced clonal shuffling between these subsets[2]. Additionally, the increase in TRS in HLA-matched naïve cells post-ICB aligns with the findings in singlet clones, suggesting the expansion of *de novo* clones driven by tumor antigen and ICB treatment.

### Carriage of HLA-matched clones is associated with improved overall survival

Given that in the single-cell data HLA-matched clones were observed to demonstrate a higher TRS, we proceeded to examine the relationship between carriage of HLA-matched clones and long-term oncological outcomes. Survival analysis across all patients receiving ICB for metastatic disease (n=231) for which we had pre-treatment samples demonstrated that patients deomstrating carriage of HLA-matched clones prior to treatment had improved overall survival (OS) (P=0.0399, log-rank test, **Supplementary Figure 11A**).

Similarly, we examined samples for which we had both pre and post-treatment data, finding that that 130/179 patients had HLA-matched clones on at least one of these two time-points. Whilst the proportion of HLA-matched clones frequently changed across patients with ICB, no patients were found to develop HLA-matched clones post-treatment who did not already have pre-treatment matched clones, with 127/130 (97.6%) patients displaying one or more HLA-matched clones across both timepoints - in keeping with the importance of the pre-treatment state for ICB activity. Notably, analysis of these patients again showed that carriage of HLA matched clones at any time point was associated with improved OS (P=0.0229,**Figure 6**), with a similar effect observed when analysis was confined to those with metastatic melanoma (P=0.0553, **Supplementary Figure 11B**).

**Figure 6.**
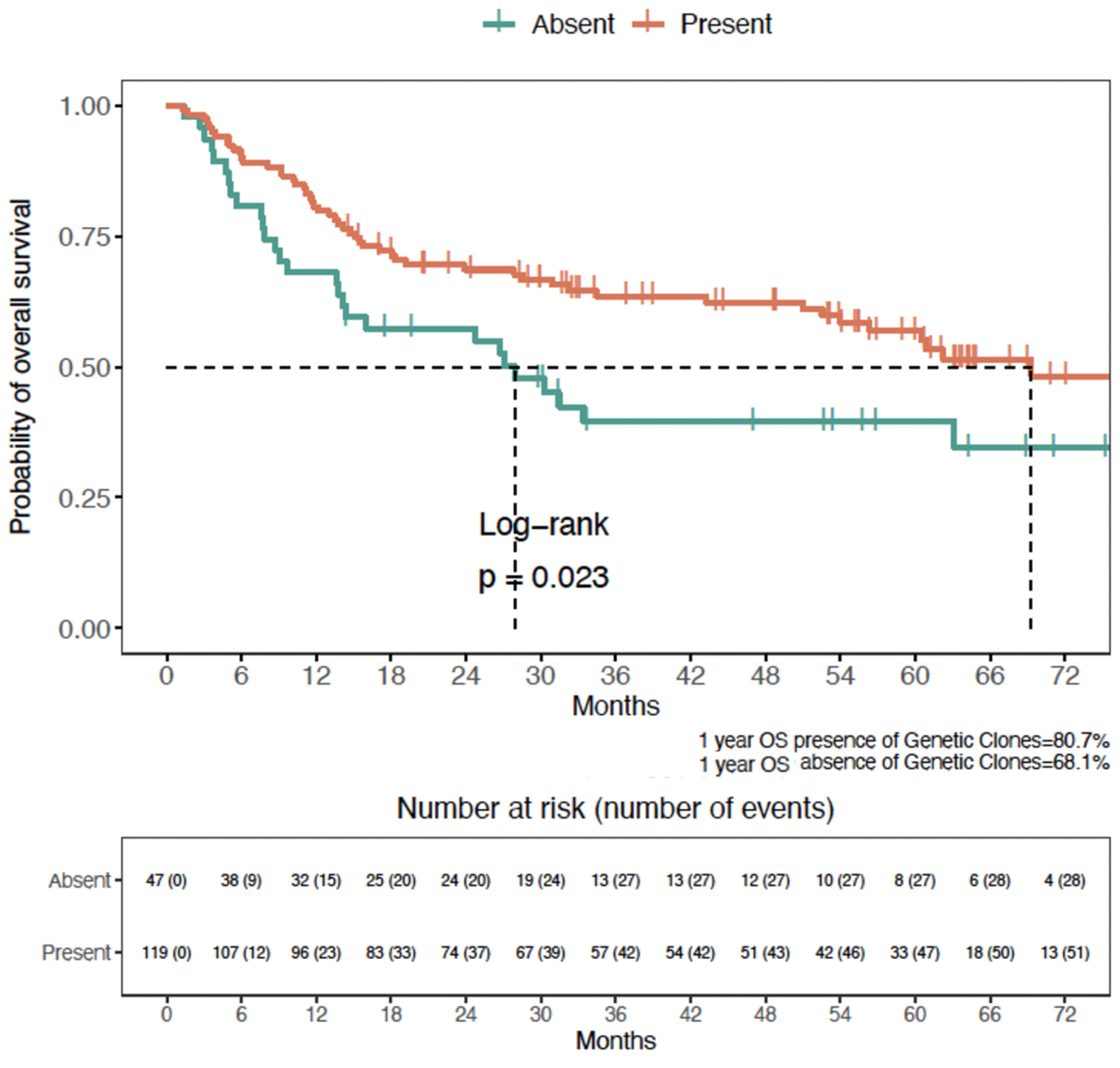
Relationship between overall survival and presence of HLA-matched clones. Kaplan Meier plot denoting that overall survival is better in patients who have a HLA-matched clone either before or after first cycle of ICB treatment.

## DISCUSSION

Here we present the first analysis of genetic determinants of CD8^+^ T cells in patients with cancer who are undergoing treatment with ICB. We describe associations both in *cis* to the V-genes of *α* and *β* TCR chains and *trans* acting associations that map to the MHC and can reduce to individual Class I HLA alleles. As well as describing the relationship between germline variation and the CD8^+^ TCR repertoire in patients with cancer, out findings support an interaction between genetic effects on the repertoire and ICB treatment with evidence that this relationship has prognostic implications.

Given our phenotype consists of unique flattened clones per individual, providing equal weighting to clones that exist as singlets and those expanded, polymorphisms associated in *cis* to the *TRAV* and *TRBV* genes likely reflect influences on V(D)J recombination propensity of clone that are selected for in that individual. Conversely, *trans* associations between MHC polymorphisms and HLA alleles with V-gene usage are proposed to reflect indirect pressures upon clonal deletion or maintenance through clonal selection via antigen presentation. In health, the drivers of this selection are both central thymic deletion of autoantigen reacting TCRs [31] and peripheral selection of antigen-reactive TCR [32]. In the context of cancer, where the burden of chronic antigen presentation leads to immune exhaustion, and also during ICB treatment, which releases immune checkpoints and favours autoimmunity causing immune related adverse events, our data suggests the major determinant of selection is successful antigen presentation to specific TCR clones. This is supported by the increased strength of association we observe for *de novo* versus persistent clones post ICB, and also the relative strength of associations in CD8^+^ TEMRA cells versus CD8^+^ Naive cells. Why non-HLA selected clones are more likely to be persistent across ICB treatment is unclear - one possibility being they’re reactive to chronic viral infection and non-exhausted. This likely speaks as to the complex processes in TCR recognition of antigen, versus the availability of specific antigens.

Our dataset is unique in that we have pre-treatment as well as post-treatment samples from the same individuals. As such, we are able to evaluate the effect of ICB on cells expressing V-genes that show association with selection to the HLA type of the individual they come from. The relatively small sample set means that only the most significant HLA:TCR V gene relationships are discernible in these data however. Nonetheless, analysis of scRNA-seq data from the same individuals demonstrates that cells carrying TCRs that show association to the HLA of the individual from which they’re obtained display gene expression profiles indicative of increased tumor reactivity compared to HLA-unmatched cells. Notably, this difference is more significant post-ICB treatment, suggesting ICB enhancement of tumor reactivity in these cells. Finally, we are able to show that patients with HLA-matched clones demonstrate increased overall survival, reflecting a direct HLA:TCR selection impact on cancer survival post-ICB.

Whilst the paired collection of samples from patients with cancer pre and post ICB means the dataset provides unique insights into the relationship between genotype, ICB treatment and TCR repertoire, we have only characterised the CD8^+^ subset. It is recognised that CD4^+^ T cells also play a role in the ICB induced anti-cancer response, especially with respect to the development of irAEs. Given the relative enrichment of Class II HLA genes with autoimmune phenotypes, future studies exploring the relationship between TCR repertoire and genotype of CD4^+^ T cells in patients with cancer will be of high importance.

In conclusion, this is the first study of the germline genetic association of TCR repertoire within cancer patients, showing significant associations at the MHC region with genes that are cancer relevant. These associations impact the survival outcomes of patients and are altered by ICB treatment.

## Acknowledgements

We are very grateful to all patients who contributed samples and participated in the study. We thank all the staff of the Day Treatment Unit, Oxford Cancer Centre, and The Brodey Centre at the Horton General Hospital. We are also grateful to all the staff of the Oxford University Hospitals NHS Foundation Trust - particularly Dr Miranda Payne, Dr Nick Coupe and Dr Rubeta Matin in the cancer centre who aid with patient recruitment, as well as the staff of the Oxford Radcliffe Biobank and Churchill Hospital Sample Handling Lab.

## Methods

### Participants

Patients were recruited from Oxford University Hospital when they were referred to receive ICB as therapy for melanoma, renal cell carcinoma and colorectal cancer. All patients provided written informed consent to donate samples to the Oxford Radcliffe Biobank (ORB) (Oxford Centre for Histopathology Research ethical approval reference 19/SC/0173, project nos. 16/A019, 18/A064, 19/A114) and allow access to their clinical data. Patients received either combined ICB (ipilimumab plus nivolumab three weekly for ≤4 treatment cycles, followed by maintenance nivolumab) or single ICB consisting of either nivolumab monthly, pembrolizumab three-weekly or pembrolizumab six-weekly. Patient demographic and clinical characteristics were collected from the electronic health record system.

### Sample collection

30-50 mL blood was collected into EDTA tubes (BD vacutainer system) just before administration of the first cycle of ICB. The blood was centrifuged to obtain peripheral blood mononuclear cells (PBMCs) and plasma by density centrifugation (Ficoll Paque). CD8^+^ T cells were isolated by CD8 positive selection using magnetic separation (Miltenyi).

### RNA extraction

Cells were resuspended in 350 µL of RLTplus buffer supplemented with beta-mercaptoethanol or Dithio-threitol. Qiashredder (Qiagen) was used to homogenise the sample, and Allprep DNA/RNA/miRNA kit (Qiagen) was then used for DNA/RNA extraction. RNA was then eluted into 34 µL of RNase-free water with concentration quantified by Qubit and DNA was eluted into 54 µl elution buffer. Both RNA and DNA samples were subsequently stored at −80°C until sequencing and genotyping.

### Bulk RNA sequencing

RNA was thawed on ice prior to mRNA isolation using Poly(A) mRNA Magnetic Isolation Module kits (NEBNext). Up to 600ng of RNA was then used to generate dsDNA libraries using NEBNext Ultra II Directional RNA Library Prep Kits as previously described [33]. Samples were then sequenced on either an Illumina HiSeq4000 (75bp paired-end) or a NovaSeq6000 (150bp paired-end). We performed TCR analysis using the MiXCR package [34] with settings as previously described [33, 2].

### Single-cell RNA and TCR sequencing

Single-cell RNA sequencing data was acquired from two different experiments. The first utilized fresh peripheral blood mononuclear cells (PBMCs) and has been previously published [2]. The second – obtained from cryopreserved PBMCs – was processed using two separate protocols. In the first protocol, dead cells were first incubated with live:dead magnetic beads (Miltenyi dead cell removal kit) and run over a MACS LD column to remove dead cells. 60,000 live PBMCs were then loaded into the partitioning reaction and processed using library kits (10X Genomics, Pleasanton, CA) following manufacturer protocols. In the second protocol, CD8^+^ T cells were first isolated by incubating with live:dead and CD8^+^ T cell negative selection beads and run over a magnetic column. 60,000 CD8+ T cells were taken from the flow-through and processed as above.

Sequencing of single-cell libraries were performed on an Illumina NovaSeq 6000 S4 flow cell (150 base pairs paired end). For each pool there were five library samples - PBMC gene expression (GEX) libraries, CD8+ T cell GEX libraries, PBMC origin TCR libraries, PBMC origin B cell repertoire libraries, CD8^+^ T cell origin TCR libraries. Batch one of sequencing was the first four pools which were sequenced across one lane of an S4 flow cell. This was deliberately under-sequenced in order to establish sequencing requirements across the rest of the samples. Following this sequencing run, there were a median of 18,000 reads/cell for GEX libraries (target 27,500) and 4,800 reads per cell for V(D)J libraries (target 7,000). We therefore sequenced the remaining seven pools plus additional sequencing of libraries from the first four pools that were under-sequenced, across three further lanes of an S4 flow cell. Across both runs we achieved a median of 28,500 reads/cell for the GEX libraries, and 7100 reads/cell for the TCR libraries.

Single-cell sequencing data were aligned against the GRCh38 human reference genome using Cellranger (v6.0.1) for GEX libraries and Cellranger VDJ for V(D)J libraries. For samples which had been sequenced across multiple lanes, the FASTQ files were inputted for alignment on the same Cellranger run. EmptyDrops [35] was used to identify empty droplets (false discovery rate (FDR) 0.01). Barcodes were excluded only if they were called as empty by both Cellranger and EmptyDrops. CellSNP-lite [36] was used to reconstruct genotype data from reads and then Vireo was used to compare this to patient-level genotype in order to de-multiplex samples from pools. Doublet identification was performed during sub-clustering by identification of mixed transcriptomes of canonical markers. Cells with <300 transcripts and >20% of mitochondrial-encoded genes were removed.

### Single-cell sequencing integration and annotation

scVI [37] was used for the integration of the two single cell experiments. We first selected 4000 highly variable genes, from the whole PBMC dataset in Scanpy [38]. An scVI model was then trained using the pool label as the batch variable and the following model parameters; number of latent variables = 30, and batch size = 1024. Once the core model was trained, datasets containing enriched cell types or higher mitochondrial proportions (10-20%) were further referenced mapped using scArches [39]. Next, clustering was implemented on a nearest neighbour graph using the 30 latent dimensions that were obtained from the scVI and scArches output. Here, the number of neighbours was set to k=30 and distance metric set to ‘cosine’. We then performed coarse leiden clustering on the graph with resolution r=0.03. For each of the resulting level 1 clusters, we calculated a new neighbour graph using scVI’s 30 latent dimensions, with the number of neighbours again set to k=30. Based on the new neighbour graph, each cluster was clustered into smaller ‘level 2’ clusters with leiden clustering at resolution r=0.3. Level 2 clusters, were then annotated based on differentially expressed genes into broad cell type categories: “CD8NK”, “CD4”, “Bcells” and “Myeloid Platelets”.

To fully capture cell type heterogeneity, we remodelled each broad ‘level 2’ subset. Per ‘level 2’ cell type, we calculated 4000 highly variable genes, re-trained scVI models using the same parameters and calculated new neighbour graphs. To guide clustering, we first performed automated cell-type annotations on scVI embeddings with Celltypist [40]. Cluster-specific marker genes were identified by performing differential expression analysis in scanpy on a given cell type compared to the rest of the cells from the same level annotation. Where clusters were highly similar, we merged them based on hierarchical dendrograms generated from gene expressions. For final CD8+ T cell annotations, cross-validation was performed based on assigned annotations from the published dataset [2].

### Quality control (QC) and HLA imputation

Genotyping was performed on the Illumina Global Screening Array 24 v3 (Illumina). For our sample QC, we used the following thresholds - heterozygosity outside 3.5 standard deviations, missing > 10% of SNPs, identity-by-descent pihat of > 0.25 and being ancestry outlier (**Supplementary Figure 3**). A total of 250 individuals were included including 231 patients with melanoma, 17 with renal cell cancer and two with colorectal cancer (**Supplementary Table 1**). For SNP QC, we used the following thresholds - MAF of 0.01, Hardy Weinberg equilibrium 0.000001 and present in at least 90% of individuals. There were a total of 486,469 SNPs tested for the GWAS. For HLA imputation, we used the Michigan Imputation Server version 2 multi-ethnic panel resulting in a total of 2,212 amino acids, 265 classical HLA alleles and 17,952 SNPs imputed. The same QC thresholds were applied to the HLA imputed data.

### Association between TCR and genetic germline variation

Bulk TCR sequencing data was processed using MixCR [34] and features were extracted, namely V-gene usage, clone counts and CDR3 amino acid sequences. Only productive clones were included. To obtain the V-gene usage phenotype, the number of unique clones with that V-gene were counted and then normalisation was performed by Trimmed Mean of M-values (TMM) function in edgeR (version 3.18) [41] followed by batch correction using removeBatchEffect function. After that, inverse normal rank transformation (INRT) was performed for each phenotype across the samples. A linear regression model was fitted in Plink (version 2.0) [42] to test each V-gene phenotype against genome wide SNPs, HLA classical alleles, amino acids/SNPs in the HLA region. We included two genetic principal components (PCs), two TCR PCs, age, gender and cancer type as covariates.

V-gene usage ⇠ genetic variant + 2 genetic PCs+ 2 TCR PCs + age + gender + cancer type

### Permutation analysis

To determine an appropriate P-value threshold for significance, we permuted it 1000 times scrambling the phenotype while keeping the same V-gene within each individual. The 5% percentile of p value was then chosen as the significance threshold. In addition to testing each HLA allele separately, we also conducted an omnibus test for each amino acid position where there are more than 1 amino acid present in the samples to investigate whether there are particular amino acid positions which are more associated to Vgenes more strongly than classical HLA alleles.

### K-mer analysis

To analyse the CDR3 amino acid sequences, K-mers of length 7 were constructed using a sliding window approach across the sequence. We chose to use 7 amino acids because the Immune-receptor-Tyrosine-based-Activation-Motif (ITAM) that is critical for the initiation of signaling following ligand engagement has a sequence of 6-8 amino acids in its centre [43].

This approach has been used successfully in other studies on TCRs [44][45]. Only sequences between length 12 and 18 were included for this analysis. The K-mers were also batch corrected and then TMM normalised followed by INRT normalised by the same procedure as the V-gene phenotype. The same linear model was fitted against HLA alleles as with the V-gene phenotype.

### Comparing pre- and post-treatment clones

Focusing on the 179 individuals for which paired samples (before and after first cycle of ICB) were available, we sub-divided clones into three groups: those only observed pre-treatment (*Unstable*), those observed post-treatment only (*Novel*), and those present across both samples (*Persistent*).

Using V-genes significantly associated with variants in the MHC region, we extracted the top classical HLA allele associated with each V-gene. HLA-matched clones were defined as clones which had this classical HLA-V-gene pairing. HLA-unmatched cells were defined as all other clones. We then calculated the proportion of HLA-matched clones for each individual within each clone group (Unstable, Persistent and Novel).

To investigate whether ICB alters the proportion of HLA-matched clones, we performed a paired Wilcoxon test on proportions of HLA-matched clones between unstable and novel groups.

For each of these three groups, we performed a V-gene vs HLA association analysis as described above for the significantly associated V-gene HLA pair.

### Single-cell TCR sequencing analysis

There were a total of 59 individuals for which we had this data as well as genotyping. Cells were included only if they had either 1 *α* and 1 *β* chain or 2 *α* and 1 *β* chain. Clones were collapsed to include only counts of unique clones and then normalised by total number of clones for the cell type being studied using the same method as bulk data. A variable representing experimental protocol was constructed. Because the experimental protocol was highly correlated with the TCR PCs, experimental protocol was regressed from the first two PCs and the residuals extracted. Then a linear model was fitted between V-gene usage and HLA alleles with age, sex, cancer type, two genetic PCs, the residuals of two TCR PCs and experimental protocol being the covariates. This was done individually for each cell type.

Single-cell gene expression sequencing was processed using Scanpy V1.10 [38] for the purpose of calculating TRS. Cells from each experimental protocol were normalised, log transformed and then scaled. To calculate tumor reactivity score for each cell, we took the sum of the 20 genes [30] and scaled it to have mean of 0 and standard deviation of 1. We then divided the cells into 2 groups, those that were HLA selected and those that were not. Using V-genes significantly associated with variants in the MHC region, we extracted the top classical HLA allele associated with each gene. HLA-matched cells were defined as cells which had this classical HLA-V-gene pairing. HLA-matched cells were defined as all other cells. To determine whether TRS was higher or lower in HLA matched cells, we performed regression of TRS score against HLA-matching status with experimental protocol as random effect and did an anova test with the null model excluding HLA-matching status. Further to that, we did this analysis separately in unique cells and cells belonging to clones (cells having the same TRA and TRB CDR3).

H1: TRS ⇠ HLA-matching status + experimental protocol H0: TRS ⇠ experimental protocol

