## Supplementary Figures for "Defining the genetic determinants of CD8^+^ T cell receptor repertoire in the context of immune checkpoint blockade"

**Supplementary Figure 1. Principal components (PC) analysis of the correlation matrix of V-gene usage** for (**A**) α chain and (**B**) β chain. This demonstrates the first PC of the α chain is dominated by TRAV1-2 which forms the key TCR for MAIT cells. Correspondingly, the PCs of the β chain correlations reflected this pattern with the first PC dominated by TRBV6-4, the key partner of TRAV1-2.

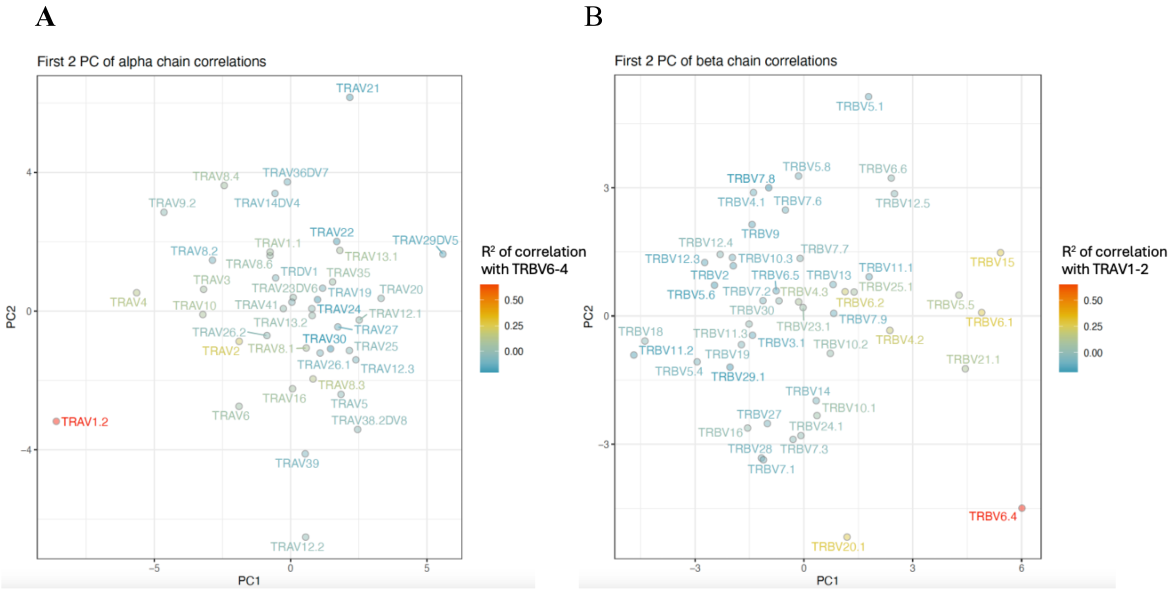


**Supplementary Figure 2. Principal components of V-gene usage of pre- and post-ICB treatment samples**. There is no clear separation between the two groups for (**A**) α chain (**B**) β chain.

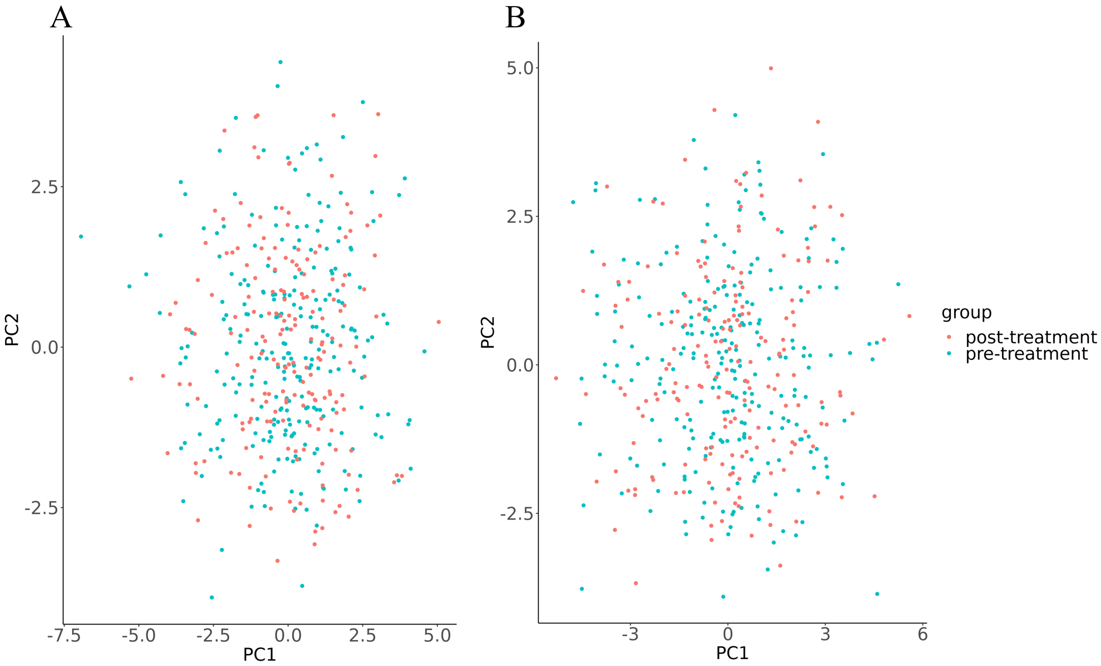

**Supplementary Figure 3. Principal components of our cohort's genetic data plotted together with the 1000 Genomes project samples** demonstrating that our samples are of European ancestry. Each of the dots represent one individual and are coloured by their population labels. AFR – African, AMR – Ad Mixed American, EAS – East Asian, EUR – European, SAS – South Asian.

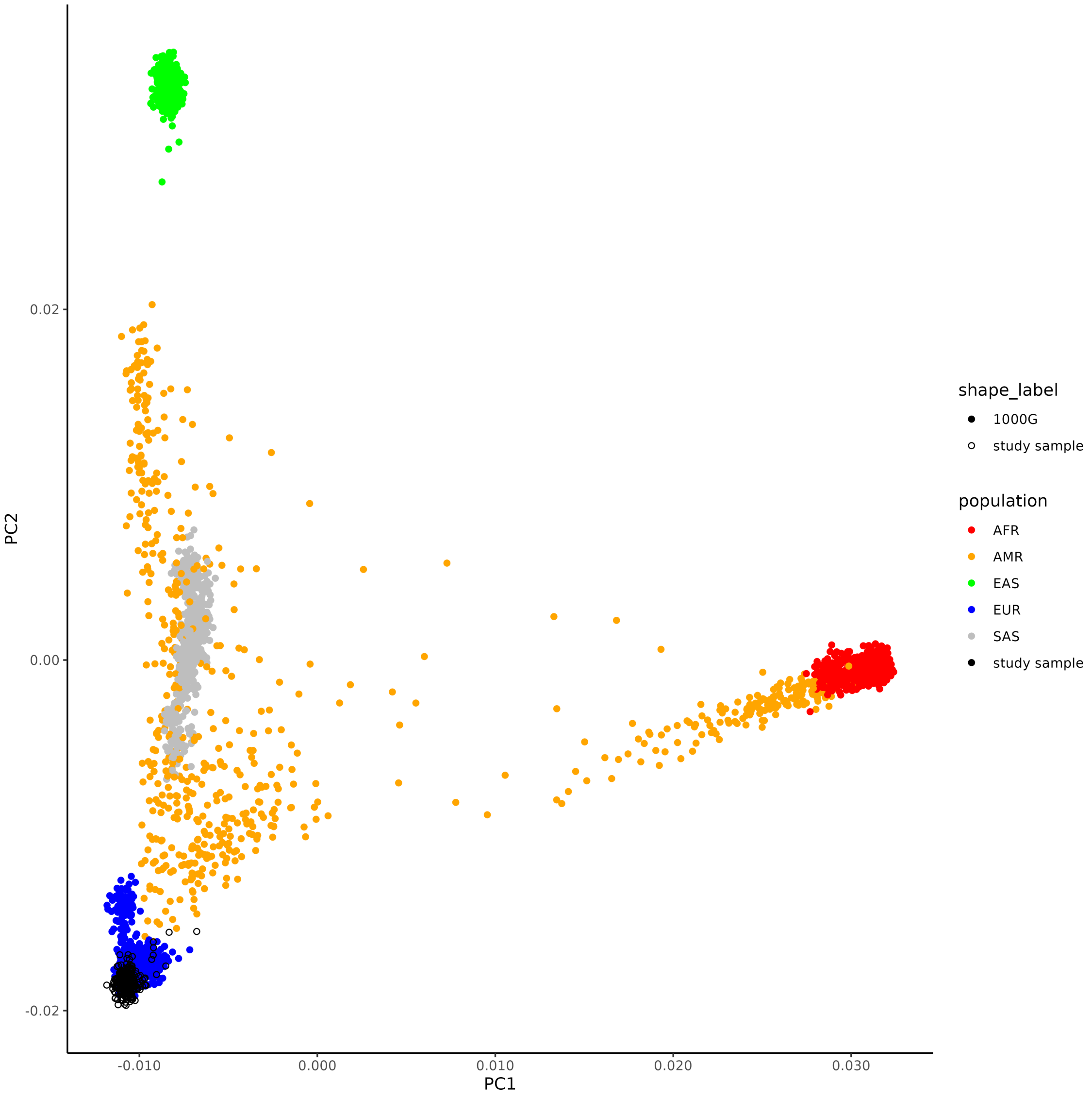

**Supplementary Figure 4. Principal components of V-gene usage** for TCR (**A**) α chain (**B**) β chain. Each point represents an individual and points are coloured by age, cancer type and gender from top to bottom respectively.

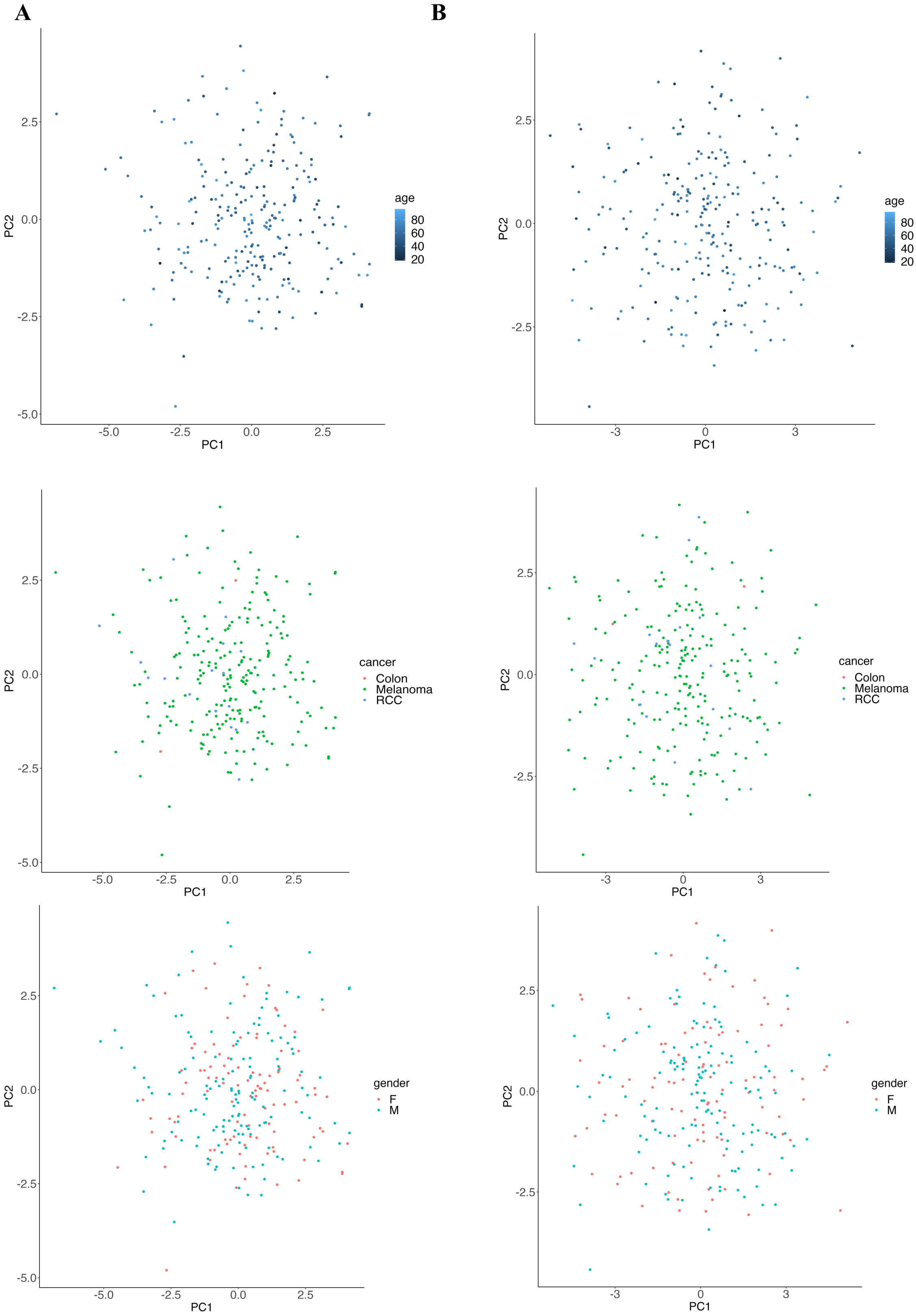

**Supplementary Figure 5. Permutation analysis.** To account for multiple testing and correlation between V-gene usage, we permuted the phenotype dataset 1000 times, preserving the V-genes that each individual had but reshuffling the samples. Plots are histograms of the permutation p-values for (**A**) α- and (**B**) β-chain V-gene usage GWAS (**C**) α- and (**D**) β chain MHC-wide association study. The 5% significance threshold is indicated by a red line.

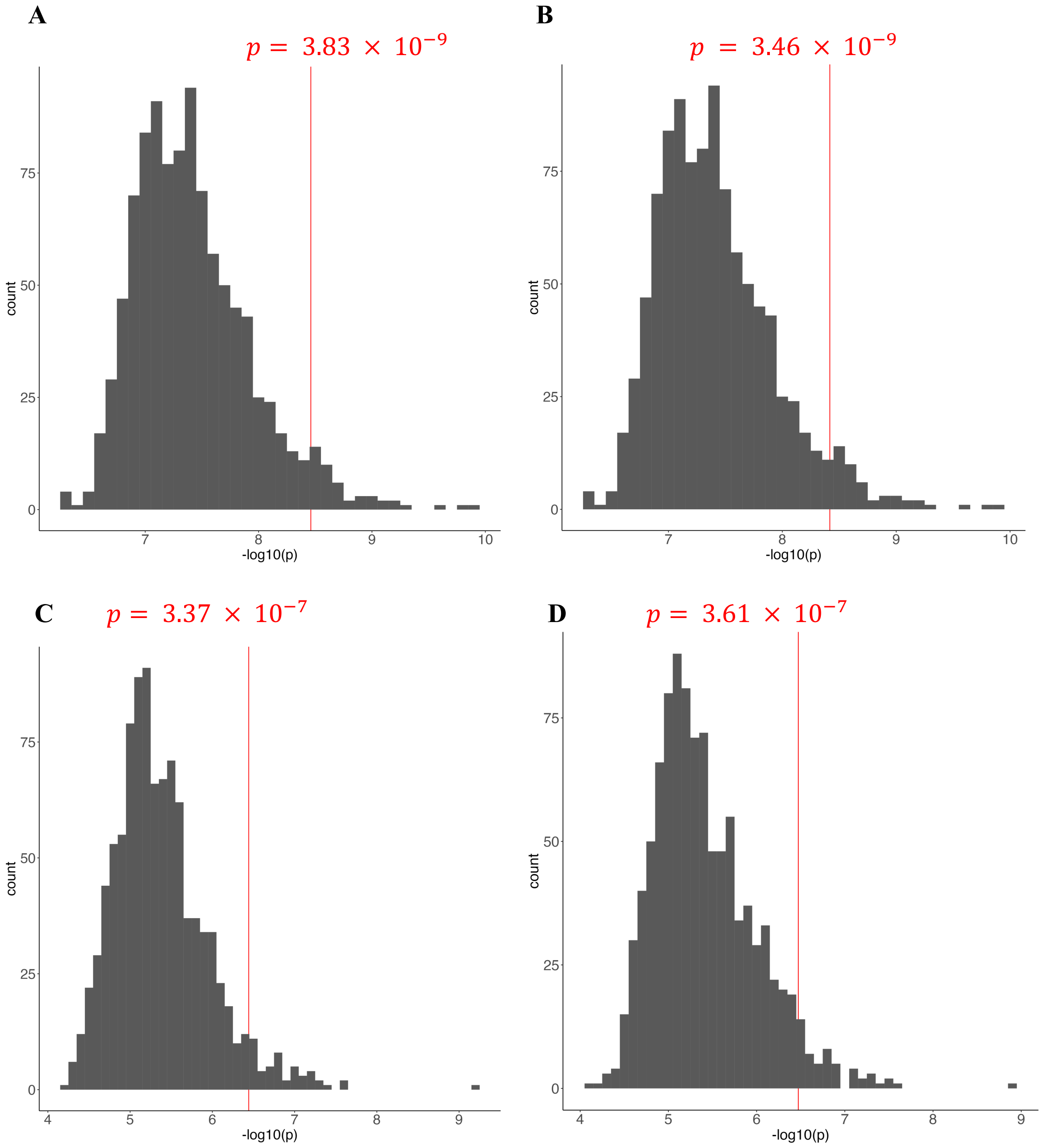

**Supplementary Figure 6. Individual locus plots of V-gene HLA associations** which pass significance threshold determined by permutation tests. Classical HLA genes are annotated in different colours annotated in the legend.

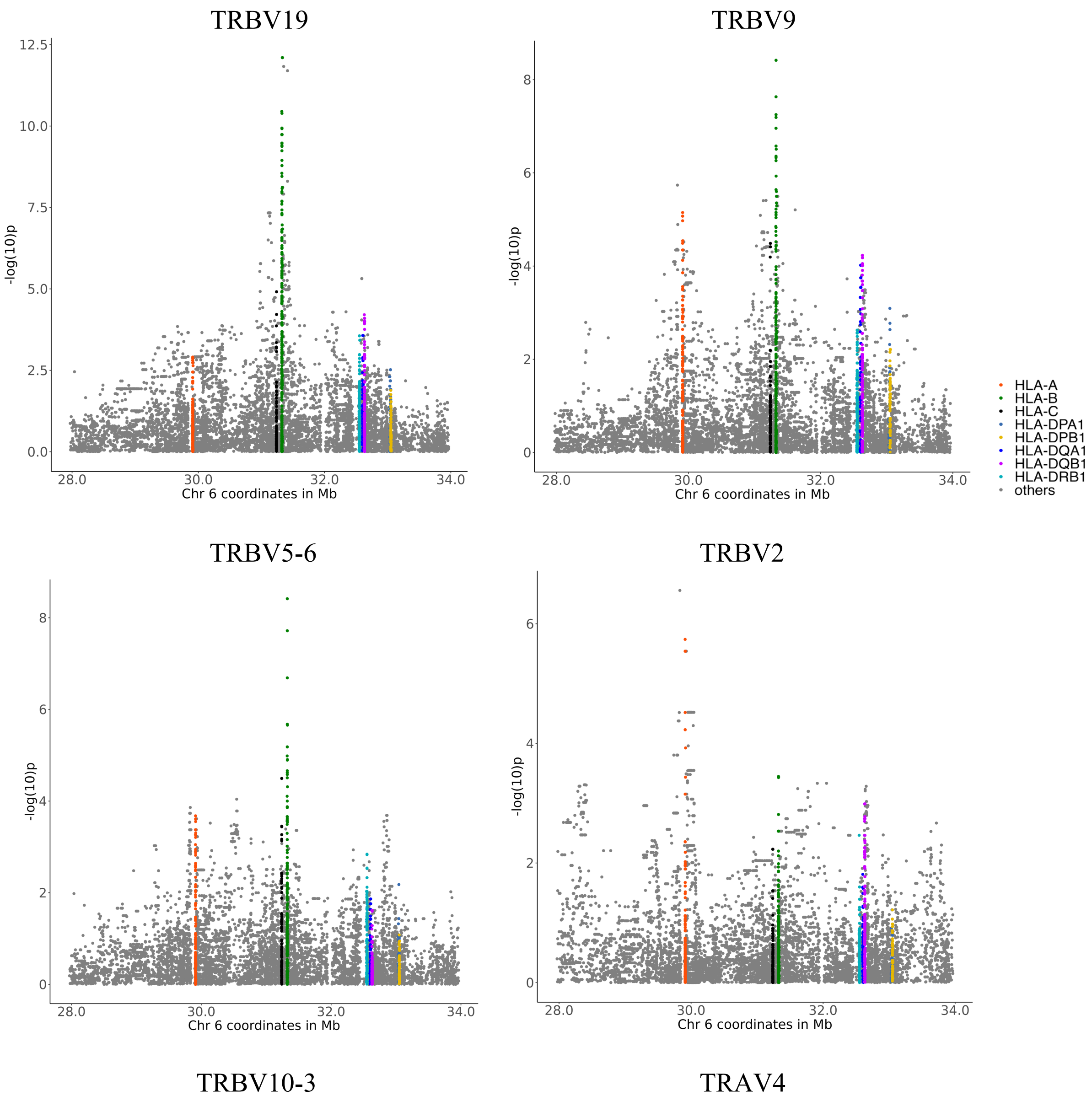

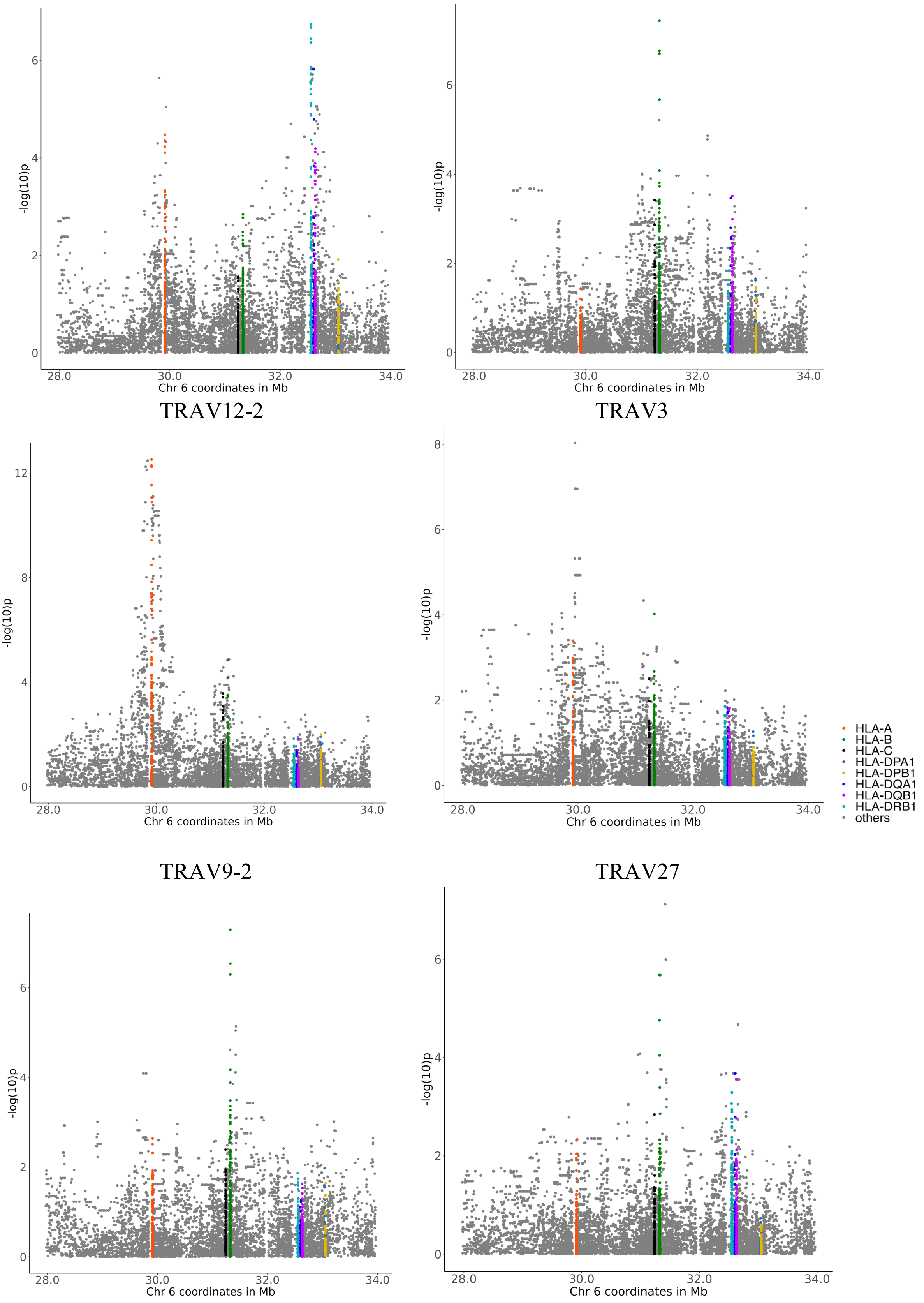

TRAV12-2
TRAV3
TRAV9-2
TRAV27
-log(10)p
Chr 6 coordinates in Mb
28.0
30.0
32.0
34.0
HLA-A
HLA-B
HLA-C
HLA-DPA1
HLA-DPB1
HLA-DQA1
HLA-DQB1
HLA-DRB1
others

**Supplementary Figure 7. Grid plots summarising the relationship between V-gene usage and 4-digit HLA**. (**A**) α chain (**B**) β chain. Each square represents the p-value of the top independent signal for each V-gene.

**A**

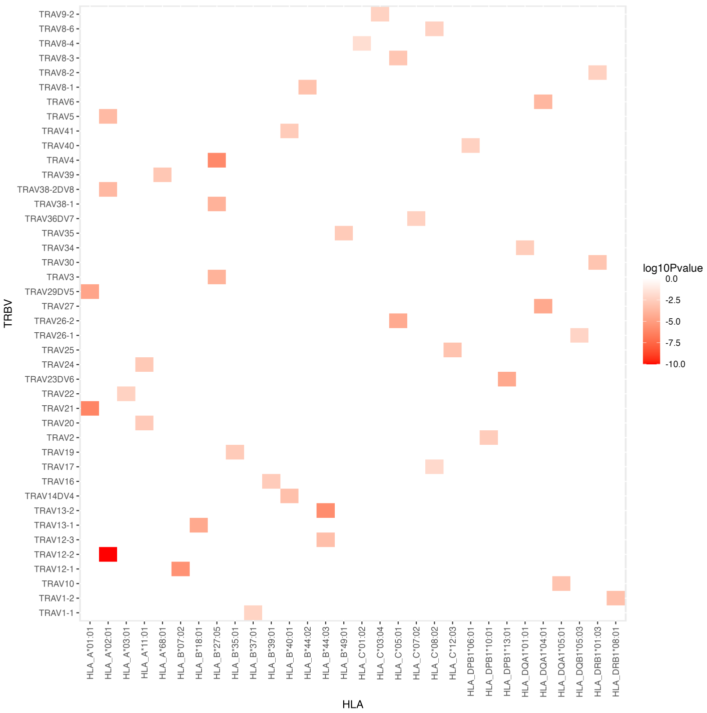


**B**

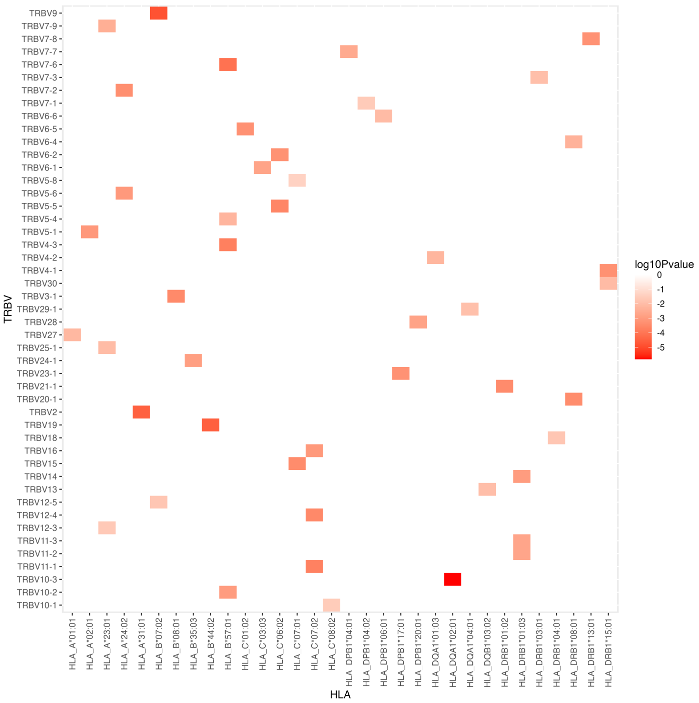

**Supplementary Figure 8. Variance explained by each of the V-genes** in the linear regression model against HLA for (**A**) α chain (**B**) β chain.

**A**

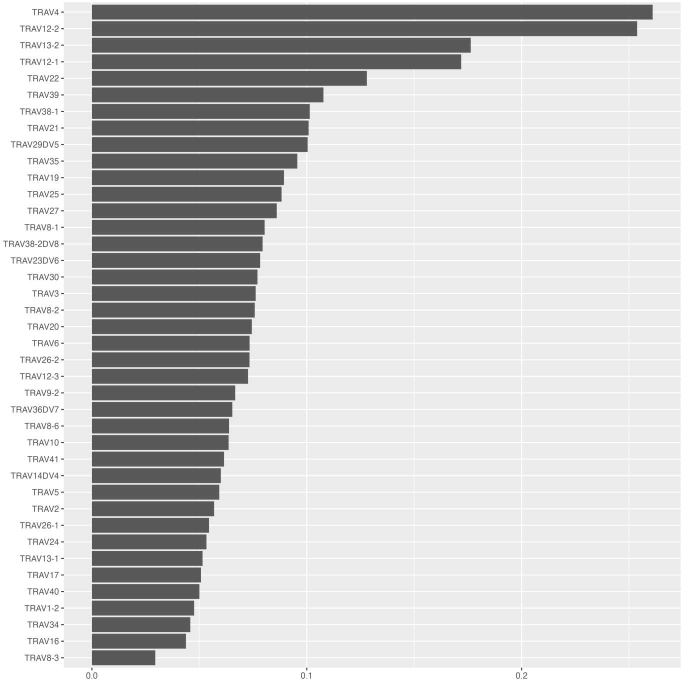


**B**

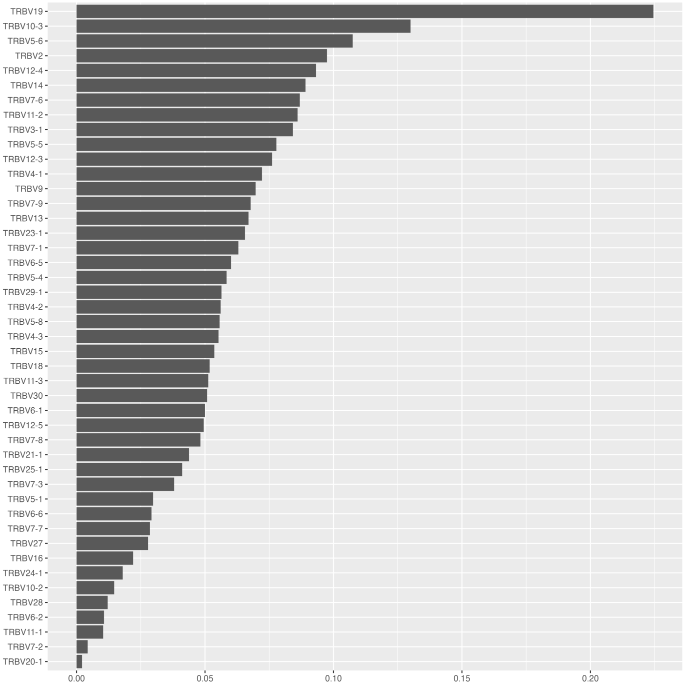

**Supplementary Figure 9. Grid plots summarising the relationship between V-gene usage and amino acid positions** in (**A**) α chain (**B**) β chain. The x-axis represents HLA and amino acid position eg. A105 is the 105th amino acid position in HLA. Each square represents the p-value of the top independent signal for each TRV gene

**A**

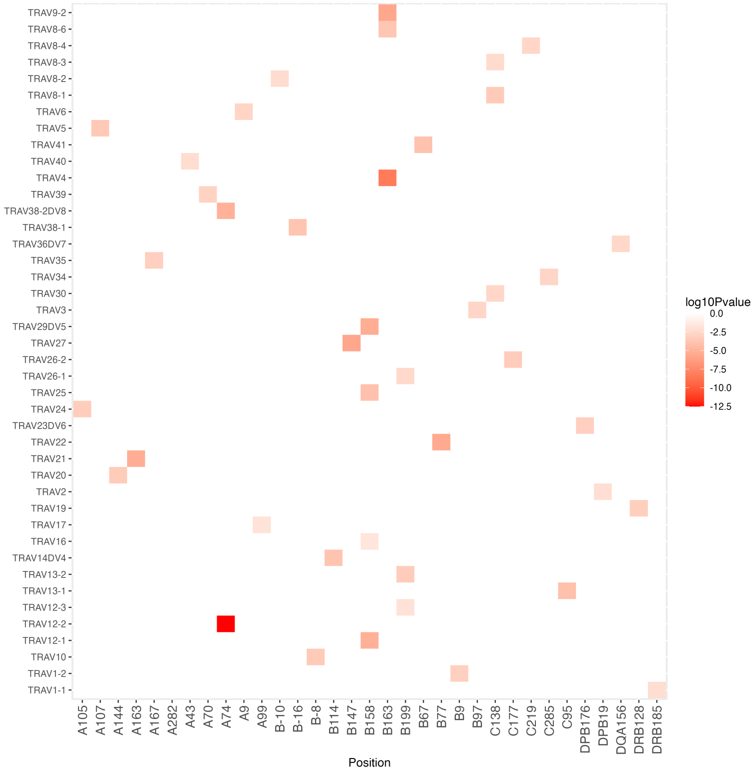


**B**

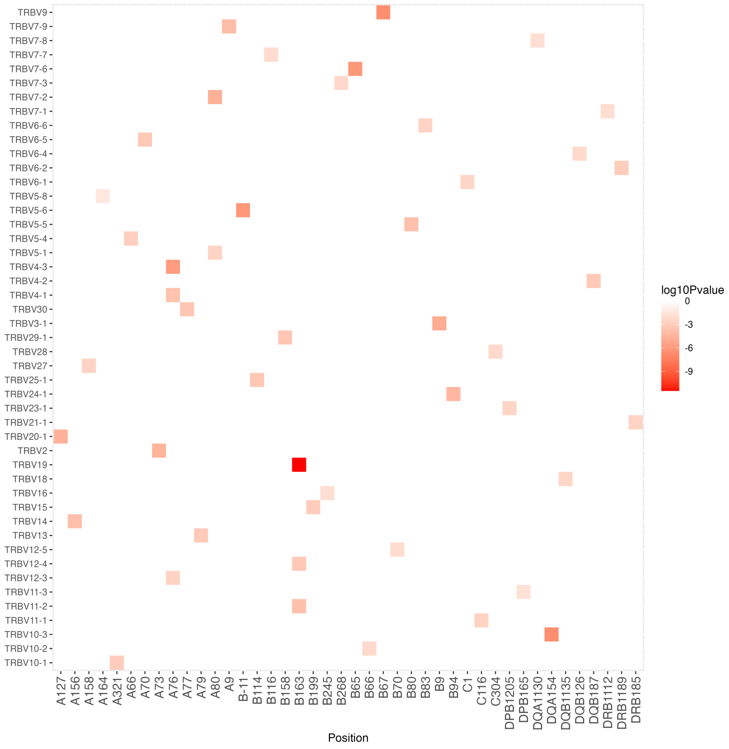

**Supplementary Figure 10. Association between TCR CDR3 K-mer usage and germline genetic variation.** Locus plot of association between variants in the MHC region and (**A**) TGDSNQP (**B**) TSGDYNE, both of which are on the TCR β chain.

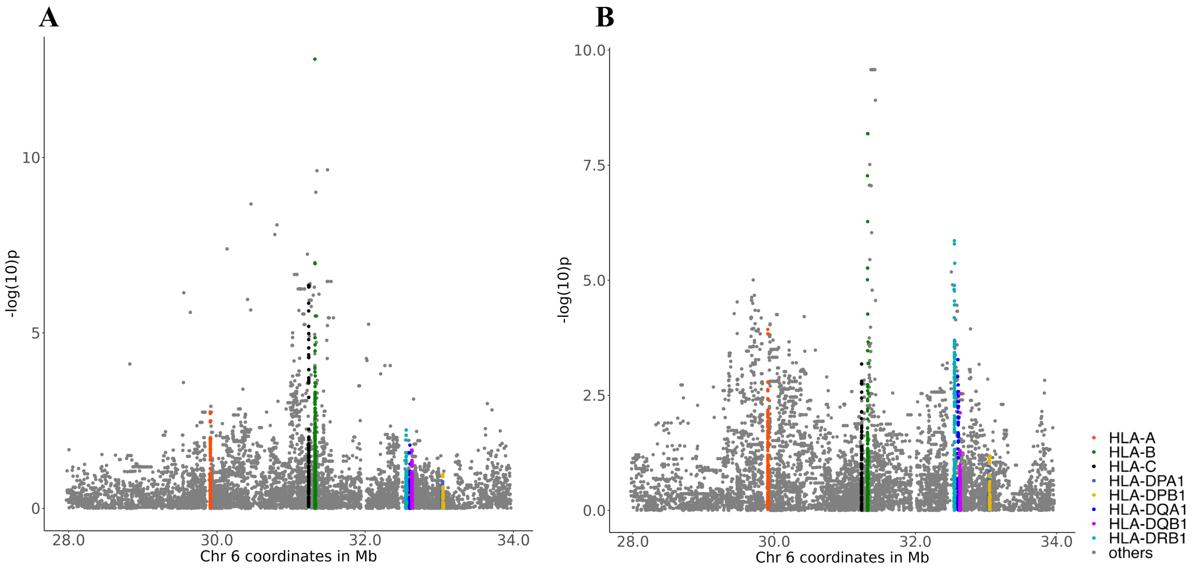


**Supplementary Figure 11. Survival analysis** of (**A**) all patients receiving ICB for metastatic disease for which we had pre-treatment samples demonstrated that patients carrying HLA-matched clones prior to treatment had improved overall survival. (**B**) patients receiving ICB for metastatic melanoma showed that carriage of HLA-matched clones either before or after treatment improved survival

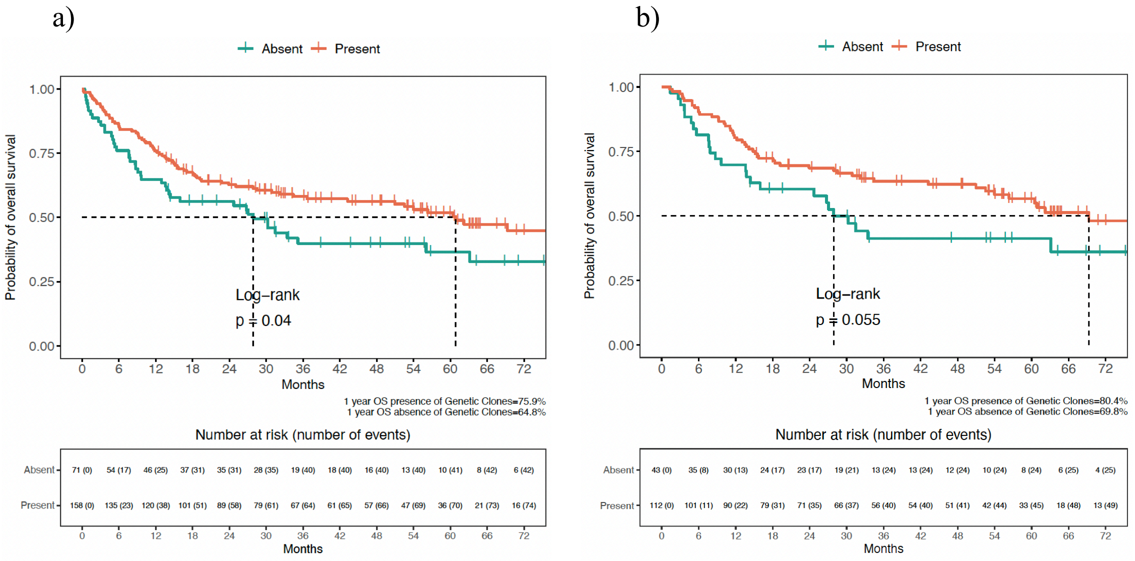
